# Application of an integrated computational antibody engineering platform to design SARS-CoV-2 neutralizers

**DOI:** 10.1101/2021.03.23.436613

**Authors:** Saleh Riahi, Jae Hyeon Lee, Shuai Wei, Robert Cost, Alessandro Masiero, Catherine Prades, Reza Olfati-Saber, Maria Wendt, Anna Park, Yu Qiu, Yanfeng Zhou

## Abstract

As the COVID-19 pandemic continues to spread, hundreds of new initiatives including studies on existing medicines are running to fight the disease. To deliver a potentially immediate and lasting treatment to current and emerging SARS-CoV-2 variants, new collaborations and ways of sharing are required to create as many paths forward as possible. Here we leverage our expertise in computational antibody engineering to rationally design/optimize three previously reported SARS-CoV neutralizing antibodies and share our proposal towards anti-SARS-CoV-2 biologics therapeutics. SARS-CoV neutralizing antibodies, m396, 80R, and CR-3022 were chosen as templates due to their diversified epitopes and confirmed neutralization potency against SARS. Structures of variable fragment (Fv) in complex with receptor binding domain (RBD) from SARS-CoV or SARS-CoV2 were subjected to our established in silico antibody engineering platform to improve their binding affinity to SARS-CoV2 and developability profiles. The selected top mutations were ensembled into a focused library for each antibody for further screening. In addition, we convert the selected binders with different epitopes into the trispecific format, aiming to increase potency and to prevent mutational escape. Lastly, to avoid antibody induced virus activation or enhancement, we applied NNAS and DQ mutations to the Fc region to eliminate effector functions and extend half-life.

## Introduction

COVID-19 cases continue to climb rapidly after causing over 80 million infections and 1.7 million deaths within a year. The causing virus, SARS-CoV-2, is identified to enter human cells by binding to the angiotensin-converting enzyme 2 (ACE2) protein, following a similar path as SARS-CoV infection in 2003 [1-3]. However, compared to SARS, mutations in the RBD domain in SARS-CoV-2 produce a stronger binding affinity to human ACE2 [4-7].

Due to the function of mediating cell entry, the spike protein and its RBD have been the focus of drug discovery for SARS coronaviruses. To date, hundreds of new research projects are focused on exploring potential treatments, many are at the preclinical trial phase, and several have reached the administration stage. For instance, the mRNA-based vaccines developed by Moderna and Pfizer-BioNTech along with the Oxford-AstraZeneca’s vaccine built on the chimpanzee adenoviral vector supplemented by the SARS-CoV-2 spike protein have been authorized for emergency use. Besides vaccines, therapeutic antibodies offer additional advantages including tractable efficacy, stability, and biocompatibility. Several antibody-based therapeutics to combat SAR-CoV-2 have been developed, including Regeneron’s REGN-CoV2 and Eli Lilly’s LY-CoV555. The former is a cocktail of two monoclonal antibodies (mAbs), REGN10933 and REGN10987, that target different RBD regions in order to maintain its neutralizing activity against future mutations [8], while the latter is isolated from a recovering COVID-19 patient [9].

While developments of new vaccines and therapeutics have progressed rapidly, SARS-CoV-2 is evolving fast pace, if not faster, and thus poses risks and uncertainties to developed candidates and products. Several variants including K417N, E484K and N501Y mutations and deletions at positions 69—70 of the RBD have been reported. One of the spike protein mutations, E484K, was suggested to hinder the neutralization effects of some polyclonal and monoclonal antibodies [10, 11]. Some early studies suggest the mRNA-based vaccines developed by Moderna and Pfizer-BioNTech may be less effective against the recently emerged South Africa variant [12, 13]. To increase neutralization likelihood and prevent mutational escape, application of a mixture of monoclonal antibodies, i.e. an antibody cocktail, results in stronger responses that are particularly effective against highly evolving pathogens [8]. Multi-specific antibody engineering based on a combination of broadly neutralizing antibodies is another highly effective method to target constantly evolving viruses. This design rationale was used to generate a trispecific antibody against HIV [14]. The underlying hypothesis is that targeting different regions of the antigen prevents resistance and escape and further enhances cross reactivity. Similar strategy using tandem linked single domain camelid antibodies showed significant efficacy against both influenza A and B viruses [15].

Several neutralizing mAbs targeting the spike RBD on the SARS-CoV virus were previously isolated and structurally characterized. Among them, the antibody 80R binds to an epitope on the RBD that largely overlaps with the ACE2 interface (Figure 1A), and a strong salt bridge is characterized as the principal component of 80R efficacy against SARS-CoV [16]. Another antibody, m396, was reported with the unique ability of blocking both virus fusion and cell entry via the spike glycoprotein [17], with its epitope overlapping with the ACE2 binding site but substantially different from the 80R’s epitope (Figure 1A). Four CDR loops, H1—H3 and L3, mediate extensive interactions with the RBD and promote strong affinity of m396 to the virus [18]. While 80R and m396 directly block the ACE2 binding site, CR3022 possess an epitope not overlapping with the ACE2 binding site (Figure 1A), making its combination with other antibodies an attractive neutralizing agent against SARS-CoV. Moreover, CR3022 was found effective against the CR3014 escape viruses and in combination with CR3014 provides prophylaxis against SARS-CoV. For instance, mutations in the SARS-CoV RBD, such as N479S and P462L, did not eliminate CR3022 neutralization potency [19]. Previous investigations reported that only CR3022 has detectable binding to the SARS-CoV-2 RBD region [20]. P384A mutation in the SAR-CoV-2 RBD was able to return the binding affinity to SARS-CoV levels which suggests that this location plays a vital role in CR3022 neutralization activity. These observations highlight the importance of optimizing the properties of these mAbs to be used for therapeutic or prophylactic purposes against SARS-CoV-2 virus.

**Figure 1.**
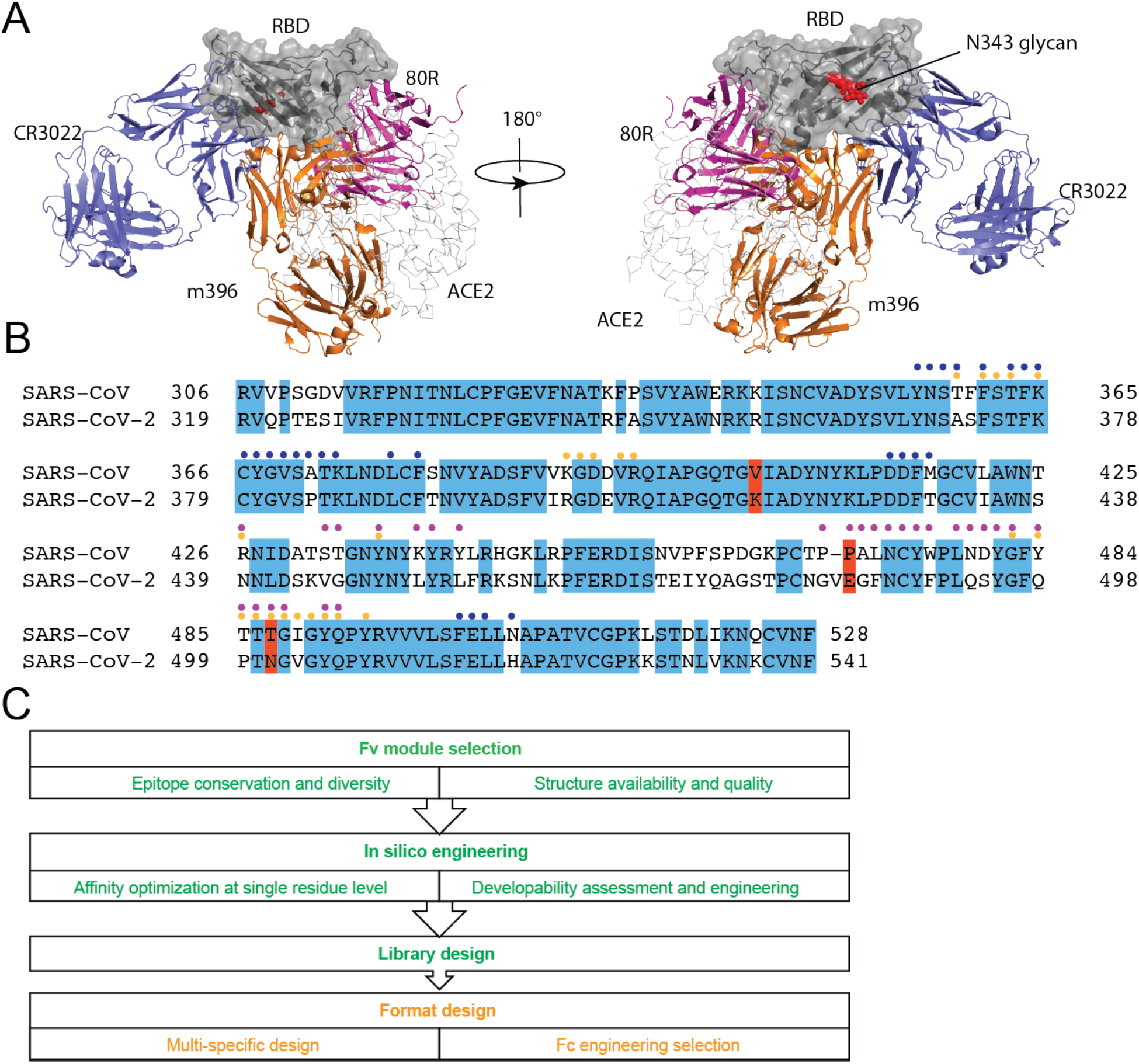
Redesign of the three anti-SARS-CoV RBD antibodies to target SARS-CoV-2. **(A)** Structural superimposition of CR3022 (cartoon representation colored in blue, PDB code 6W41), m396 (cartoon representation colored in orange, PDB code 2DD8), 80R (cartoon representation colored in magenta, PDB code 2GHW), and ACE2 (ribbon representation colored in grey, PDB code 6M17) on their binding to SARS-CoV or SARS-CoV-2 RBD. N-glycosylation at N343 site is shown as red sphere, while glycosylation at N313 site is not visible in the crystal structures. **(B)** Sequence alignment of the SARS-CoV-2 and SARS-CoV RBDs. Conserved residues between SARS-CoV and SARS-CoV-2 are highlighted in blue color. Recent UK and South African SARS-CoV-2 mutation sites are highlighted in red. Epitope residues are indicated by colored dots: blue for CR3022, orange for m396, and magenta for 80R. **(C)** Schematic workflow for engineering of the three antibodies. Green text indicates engineering toward developability and cross reactivity, and orange text indicates format related designs in Fab and Fc regions.

Discovery of antibody therapeutics has rapidly evolved in the past few years, and research in lead generation and optimization faces strong challenges in needing high success rates and short timelines. Structure-based rational engineering of antibodies has been shown fast and highly effective in optimizing features of lead candidates, including cross reactivity, potency, developability, and safety profile. Hereto we selected the above mentioned three structurally known anti-SARS-CoV monoclonal antibodies with established neutralization potency and fed them into our computational design pipeline to propose SARS-CoV-2 neutralizing antibodies. Moreover, combinations of those binders are designed into a multi-specific format aiming to further enhance the anti-viral potency and tolerance to viral evolution in the RBD.

## Method

### Selection of templates

SARS-CoV and SARS-CoV-2 share the same RBD-ACE2 interface as a cell entry path. The RBDs have 76% sequence identity between SARS-CoV and SARS-CoV-2, and the level of identity decreases to 64% within the RBD-ACE2 interface residues [4] (Figure 1B). Crystal or cryoEM structures of multiple anti-SARS-CoV Fab complexes with the RBD from SARS-CoV or SARS-CoV-2 are available; we select three clones, m396, 80R, and CR3022 as our templates, with the filtering criteria of continuously overlapping epitopes, ranging from highly conserved RBD surface to more mutation prone (Figure 1A&B).

### Developability assessment and engineering at Fv level

The Fv of the candidates were isolated from their complex structure and subjected to computational prediction of developability features including surface patches, chemical degradation of Asp and Asn, and oxidation of Met. Patch calculation included spatial aggregation propensity (SAP) [21] using Discovery Studio (BIOVIA, Dassault Systèmes) with a 5 Å radius and clustering of residues in the patch analysis using Molecular Operation Environment (MOE) version 2019.0102 [22]. Patches larger than 50 Å^2^ were selected for further visual inspection. Deamidation and isomerization motifs were analyzed with bioMOE using structure-based prediction models developed by Sydow et al. [23] and Robinson et al. [24]. Risk of methionine oxidation was predicted using sulfur solvent-accessible area and 2-shell models with bioMOE [25]. Residue scanning on the patch residues or chemical liability motifs were manually inspected and mutation strategies were made following two criteria: 1) mutation does not impact binding; and 2) mutation reduces patch area.

### Structures preparation for SARS-CoV-2 reactivity engineering

All antibody sequences reported here are renumbered using continuous peptide numbering. The RBD from SARS-CoV-2 spike structure is used to replace the RBD in the m396 and 80R complexes. For 80R, the single chain Fv (scFv) was split to Fv with standard VH-VL pairing and the linker between VH and VL in the scFv was removed. Antibody residues that are within 6 Å of the RBD are selected and fed to residue scanning in MOE, Rosetta, TopNetTree, and SAAMBE3D. To prepare the structures for residue scanning, the PDB model of Fab/Fv with RBD2 were initially protonated and energy minimized with MOE. For calculations in Rosetta and machine learning based methods, the MOE minimized structure was further relaxed with Rosetta.

### MOE

The MOE computation workflow, unless specified, was performed with MOE.2019.01.02 [22] with Amber10 forcefield [26] and Born solvation model [27]. After protonation and minimization, all selected residues that are within 6 Å of the antigen were subjected to single residue scanning to 20 natural residues with ensemble LowMode [28]. For ensemble generation, residues located outside 4.5 Å away from the mutation site were fixed.

### Rosetta Flex ddG

Flex ddG is built upon the Rosetta architecture and incorporates the conformational sampling of backbone and side chain torsions into the free energy calculation using the Talaris scoring function in Rosetta[29]. Following the nonlinear reweighting protocols, i.e. generalized additive models, of the Rosetta energy function computed for each structure of mutant and wildtype at complex and unbound states, Flex ddG estimates the ΔΔG values. Firstly, the three RBD-Fv complex structures prepared by MOE were energy minimized using the Rosetta FastRelax protocol. For each complex, the lowest energy structure was chosen from the 10 relaxed structures and used for the next step. Secondly, ΔΔG estimates for each single point mutation were calculated using the “Flex ddG” protocol with default parameters as described in the reference [30], except for using 10 instead of 35 averaged models due to computational constraints. This change was made according to the observation in the original publication that the correlation and mean absolute error between predicted ΔΔG and experimental ΔΔG became stable when the number of averaged models was around 10 or more [30].

### TopNetTree

TopNetTree is a machine learning (ML) model that utilizes site-specific persistent homology to extract the local geometric information of the protein complexes and mutation sites [31]. As such, this method simplifies the complexity of the 3D atomic structure and in conjunction with ML methods, including convolutional neural networks and gradient-boosting trees, it is able to capture the change in the underlying biochemical features, such as hydrogen bonding and dispersion interaction represented at the zeroth homology group H_0_, along with the structural change, represented at first and second homology groups H_1_ and H_2,_ at the mutation site. The model is trained and validated on different single site mutation datasets, including computational and experimental data, such as SKEMPI v2.0 [32] and AB-Bind [33]. Validation results of this method illustrate satisfactory performance across different databases and mutation regions (accessible surface area) for the ΔΔG prediction. The ΔΔG calculations were performed using both topological and physiochemical properties. The original TopNetTree model parameters were used in this study. The optimized complex structures obtained from Rosetta were used as input for free energy calculations. To maintain consistency with TopNetTree methodology each structure was further optimized with the profix module in Jackal modeling suite.

### SAAMBE-3D

SAAMBE-3D is an ML based model that is constructed based on a variety of features spanning across multiple chemical, physical, sequential and mutation specific properties. This allows SAAMBE-3D to efficiently extract essential information from the structure and predict the ΔΔG upon mutation. We downloaded and used, without modification, the scripts and models associated with the publication [34] (http://compbio.clemson.edu/saambe_webserver/index3D.php). The model was trained on 3753 single point mutations from 299 different protein–protein complexes, of which approximately 650 mutations were from 76 Ag-Ab complexes. We did not re-train the model on the more relevant Ag-Ab subset as the significant reduction in the dataset size may decrease the performance of the model. Rosetta optimized structures for each Fv-RBD system were used as the initial structure for SAAMBE-3D calculations.

### Consensus Z-score

Z-scores were used to extract the favorable mutations for each system. Coupled with the structural inspection, z-scores have been shown to accurately highlight/guide mutation selection from the vast affinity maturation calculations. We used a modified Z-score as suggested by Sulea et al. [35] where the median and median absolute deviation (MAD) were used based on the following equation:

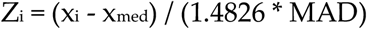

Each Z-score was averaged over the four methods. Top 60 average scores for each system supplemented with the structural inspection to select the final list of affinity promoting mutations.

## Results

### Selection of three neutralizing antibodies

Monoclonal antibodies 80R, m396, and CR3022 have been well characterized to prove their neutralizing potency to SARS-CoV virus. The mutations between SARS-CoV-2 and SARS-CoV RBD make these neutralizers not immediately applicable to block the RBD-ACE2 interactions [36, 37]. Publicly available high-quality structures of Fab in complex with RBD allow us to quickly design SARS-CoV-2 binders through our structure-based rational engineering platform, which has been serving our cross reactivity and affinity maturation engineering purposes in biologics projects [38]. The epitopes of these three antibodies are located in relatively conserved surfaces on the RBD (Figure 1A&B). The 80R and m396 epitopes largely overlap with the ACE2 binding site, which limits the possibility of escaping mutations on the RBD as mutations abolishing ACE2 interaction are unfavorable. Although the CR3022 epitope is distal from the ACE2 binding site, it has been shown as a conserved epitope between SARS-CoV and SARS-CoV-2 [20]. Additionally, the glycosylation sites (N331 and N343) in the SARS-CoV-2 RBD are away from the epitopes of the three antibodies, making it less likely to shield antibody binding (Figure 1A) [39]. Lastly, 80R and CR3022 utilize kappa, while m396 uses lambda light chain. The difference in light chains also helps assembly design into multi-specific antibodies and minimize mispairing risks.

### In silico mutagenesis and consensus Z-score

For each complex structure, antibody residues within 6 Å from the RBD were selected for ΔΔG calculations upon mutation to all 20 amino acids. This resulted in 48, 35, and 34, mutation sites corresponding to 80R, m396, and CR3022, respectively. Figures 2D, 3D, and 4D depict the results of ΔΔG calculations performed on 80R, m396, and CR3022, respectively, using the 4 computational methods discussed before. Due to the mutational structure sampling algorithms, the binding affinity scores comparing mutations to wild type (e.g. H:S101S) can be nonzero. For normalization, the ΔΔG value for each mutation is offset so that the wild type mutations are zero. Interestingly, predicted ΔΔG values obtained from SAAMBE-3D are mainly unfavorable (positive values), and the range of predicted values is smaller than other methods. Another observation is the large variation of predicted values among these four methods, reflecting the need of an approach to effectively rank the mutations. Previous studies in binding affinity predictions suggest that using a consensus approach over different methods can improve prediction accuracy [35, 40-45]. Following this rationale, we applied a similar strategy to rank the single mutations from the four computational predictions for each antibody. We used relative ranking instead of absolute score due to different magnitudes and scales of the four methods. A Z-score describes a value’s relationship to the mean of a group of values, which is useful for normalization of raw scores. Here we used a Z-score based on the median value instead of the mean value for each scoring function, which reduces the sensitivity of Z-scores to outliers. By averaging the Z-score from the four methods, consensus Z-scores were computed, and the top ranked mutations were visualized to validate the predictions. For each system we selected the top 60 mutations as presented in Table 1.

**Figure 2.**
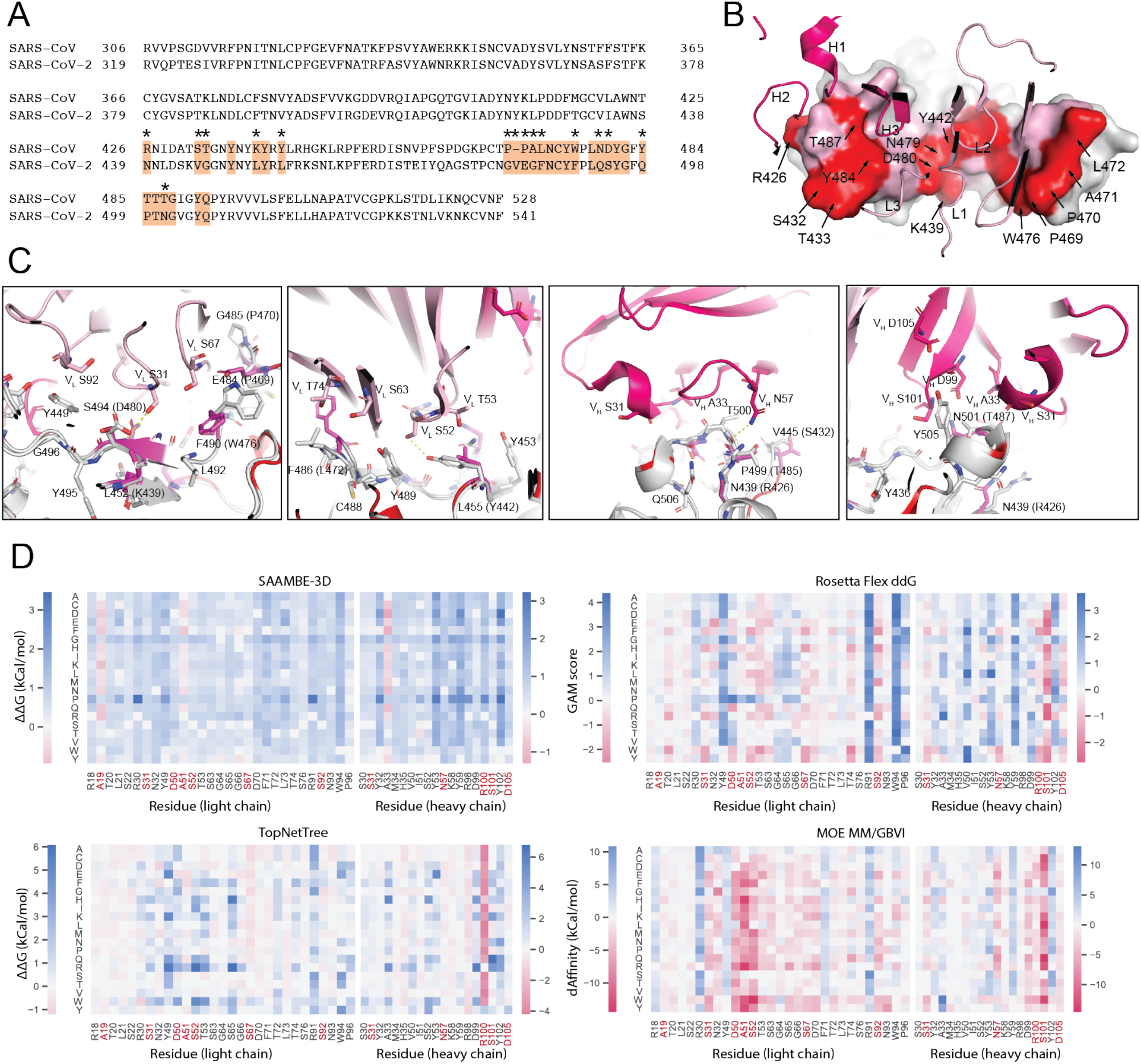
Engineering of 80R. **(A)** Sequence alignment of SARS-CoV-2 and SARS-CoV RBDs. 80R epitope residues are highlighted in orange. Non-conserved epitope residues are marked with asterisks. **(B)** Epitope residues on SARS-CoV-2 are shown. CDR loops are labeled. Epitope residues that are conserved between SARS-CoV-2 and SARS-CoV are shown in pink, and those that are not conserved are shown in red. **(C)** Interactions between selected 80R residues for engineering and epitope residues are shown. Amino acid variants observed in SARS-CoV are in parentheses. SARS-CoV-2 RBD is grey, 80R heavy chain is magenta, and 80R light chain is pink. Residues are numbered according to their positions on the SARS-CoV-2 S protein sequence. **(D)** Heatmap of prediction of all possible mutations for selected residues on 80R from SAAMBE-3D, TopNetTree, Rosetta flex ddG, and MOE MM/GBVI methods. Residues selected for library design are colored in red.

**Figure 3.**
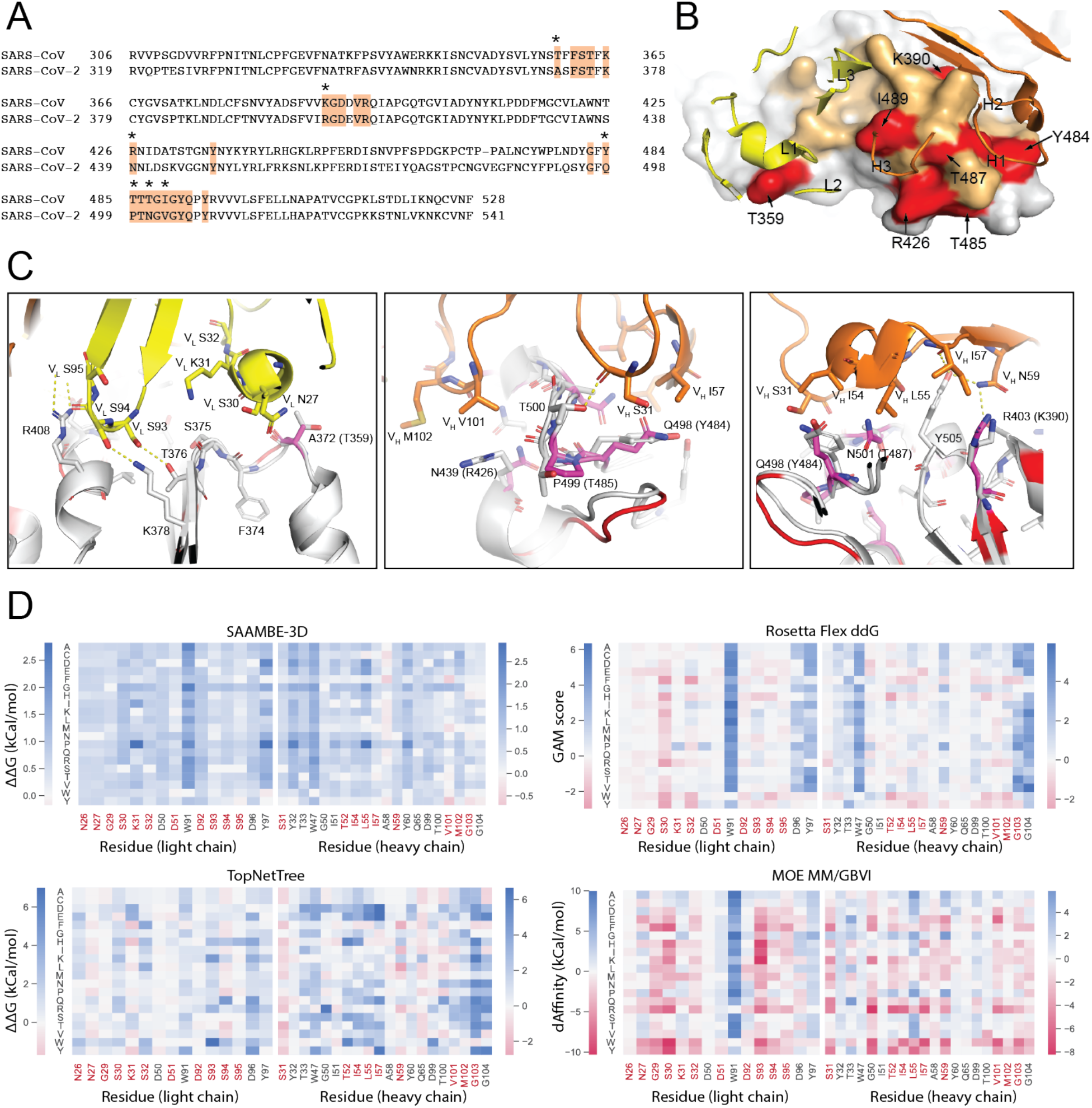
Engineering of m396. **(A)** Sequence alignment of SARS-CoV-2 and SARS-CoV RBDs. M396 epitope residues are highlighted in brown. Non-conserved epitope residues are marked with asterisks. **(B)** Epitope residues on SARS-CoV-2 are shown. CDR loops are labeled. Epitope residues that are conserved between SARS-CoV-2 and SARS-CoV are shown in orange, and those that are not conserved are shown in red. **(C)** Interactions between selected m396 residues for engineering and epitope residues are shown. Amino acid variants observed in SARS-CoV are in parentheses. SARS-CoV-2 RBD is grey, m396 heavy chain is orange, and m396 light chain is yellow. Residues are numbered according to their positions on the SARS-CoV-2 S protein sequence. **(D)** Heatmap of prediction of all possible mutations for selected residues on m396 from SAAMBE-3D, TopNetTree, Rosetta flex ddG, and MOE MM/GBVI methods. Residues selected for library design are colored in red.

**Figure 4.**
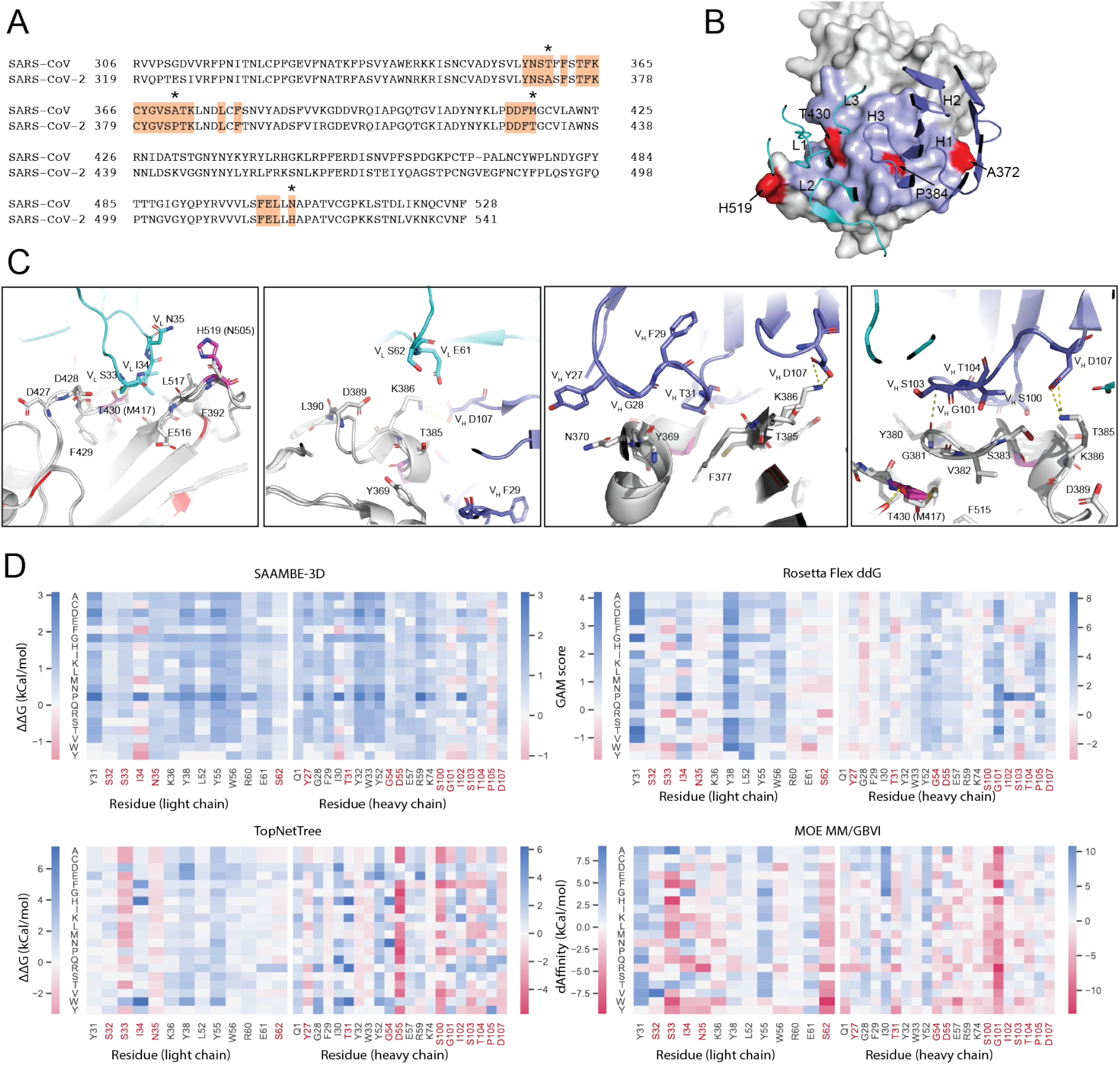
Engineering of CR3022. **(A)** Sequence alignment of SARS-CoV-2 and SARS-CoV RBDs. CR3022 epitope residues are highlighted in brown. Non-conserved epitope residues are marked with asterisks. **(B)** Epitope residues on SARS-CoV-2 are shown. CDR loops are labeled. Epitope residues that are conserved between SARS-CoV-2 and SARS-CoV are shown in blue, and those that are not conserved are shown in red. **(C)** Interactions between selected CR3022 residues for engineering and epitope residues are shown. Amino acid variants observed in SARS-CoV are in parentheses. SARS-CoV-2 RBD is grey, CR3022 heavy chain is blue, and CR3022 light chain is cyan. Residues are numbered according to their positions on the SARS-CoV-2 S protein sequence. **(D)** Heatmap of prediction of all possible mutations for selected residues on CR3022 from SAAMBE-3D, TopNetTree, Rosetta flex ddG, and MOE MM/GBVI methods. Residues selected for library design are colored in red.

**Table 1.** Top 60 mutations ranked by consensus Z-scores. Mutations are represented in “Chain ID: Mutation” format and are associated with consensus z-scores calculated by the formula in the Methods section.

| 80R |  | m396 |  | CR3022 |  |
| --- | --- | --- | --- | --- | --- |
| chain:mutation | Z score | chain:mutation | Z score | chain:mutation | Z score |
| H:S101W | -3.068 | L:S30Y | -2.865 | H:S103W | -2.238 |
| H:R100W | -3.029 | L:S30F | -2.464 | H:G101F | -2.169 |
| H:S101M | -2.905 | L:S30W | -2.351 | L:S33H | -1.939 |
| H:S101L | -2.886 | H:L55Y | -2.241 | L:S33F | -1.828 |
| L:A51F | -2.604 | H:V101Y | -1.863 | H:G101A | -1.653 |
| L:A51H | -2.558 | L:S93H | -1.797 | L:S33M | -1.630 |
| H:R100F | -2.48 | H:S31R | -1.788 | H:S103F | -1.628 |
| L:A51Y | -2.384 | L:S30R | -1.695 | L:S33Y | -1.470 |
| H:S101D | -2.292 | H:V101W | -1.664 | L:S33E | -1.466 |
| H:S101C | -2.219 | H:S31F | -1.656 | L:S62W | -1.433 |
| H:R100M | -2.114 | H:G103F | -1.637 | H:S103Y | -1.423 |
| H:R100L | -2.099 | H:S31Y | -1.545 | L:S33L | -1.316 |
| L:A51L | -2.09 | L:S30H | -1.507 | H:T104Q | -1.306 |
| L:A51Q | -1.957 | H:N59F | -1.496 | H:G101W | -1.291 |
| L:A51I | -1.931 | H:I54F | -1.459 | H:D55Q | -1.288 |
| H:R100E | -1.898 | L:S94F | -1.428 | L:N35R | -1.241 |
| H:R100I | -1.892 | H:T52Y | -1.413 | H:D55I | -1.213 |
| H:R100D | -1.887 | H:S31Q | -1.38 | H:D55H | -1.199 |
| H:R100V | -1.874 | L:S30E | -1.366 | H:S103M | -1.139 |
| H:S101I | -1.833 | L:S93E | -1.365 | L:S33Q | -1.137 |
| L:A51W | -1.824 | L:S95F | -1.356 | L:S33I | -1.131 |
| L:S67Y | -1.816 | L:S93Y | -1.339 | H:S100Q | -1.126 |
| L:D50E | -1.791 | L:S93I | -1.329 | L:S62Y | -1.114 |
| L:A51E | -1.768 | H:M102W | -1.29 | H:D55C | -1.114 |
| L:S67H | -1.754 | L:S32Y | -1.263 | L:S62V | -1.111 |
| L:S67F | -1.742 | H:V101F | -1.263 | H:T31E | -1.110 |
| H:R100C | -1.739 | H:N59R | -1.248 | H:D55S | -1.087 |
| H:S101V | -1.731 | H:S31W | -1.244 | H:I102Y | -1.085 |
| L:S52F | -1.697 | H:G103W | -1.244 | H:T104E | -1.077 |
| H:R100K | -1.688 | L:G29F | -1.24 | H:T31M | -1.069 |
| L:S92H | -1.641 | H:L55F | -1.237 | L:S62M | -1.055 |
| L:A51M | -1.626 | H:I57R | -1.213 | H:S100P | -1.055 |
| H:R100A | -1.612 | H:V101L | -1.21 | H:S100A | -1.040 |
| H:R100S | -1.556 | H:S31M | -1.208 | H:Y27R | -1.030 |
| H:R100N | -1.523 | H:T100W | -1.198 | H:S100M | -1.010 |
| L:A51K | -1.512 | L:N27E | -1.187 | H:Y27W | -0.998 |
| L:S31Y | -1.422 | H:I57W | -1.184 | H:D107W | -0.998 |
| H:R100T | -1.405 | H:N59Y | -1.178 | H:D55G | -0.960 |
| H:S101T | -1.362 | H:G50E | -1.163 | H:P105W | -0.959 |
| L:D50Y | -1.349 | H:S31K | -1.159 | H:G101S | -0.958 |
| H:R100Q | -1.336 | H:M102Y | -1.134 | L:S33W | -0.956 |
| H:S101E | -1.331 | L:S30K | -1.132 | H:S100T | -0.939 |
| L:A51D | -1.303 | L:S32W | -1.121 | L:I34Q | -0.930 |
| L:S92M | -1.29 | H:V101M | -1.11 | H:T31I | -0.923 |
| L:S67M | -1.274 | L:S93N | -1.104 | L:S62T | -0.920 |
| L:A51R | -1.273 | H:G50Q | -1.062 | H:I102F | -0.920 |
| H:D105W | -1.269 | L:G29R | -1.051 | L:S62R | -0.900 |
| L:S52M | -1.23 | L:S30M | -1.03 | L:S62N | -0.882 |
| L:S67E | -1.217 | L:G29Y | -1.013 | L:S62L | -0.858 |
| H:S101Y | -1.217 | H:I57H | -1.011 | L:S33C | -0.832 |
| H:S31W | -1.199 | L:S94W | -0.993 | L:S62F | -0.809 |
| L:S67K | -1.187 | H:V101R | -0.976 | L:N35Y | -0.800 |
| L:S92R | -1.172 | L:S95I | -0.96 | L:S62H | -0.789 |
| L:A19K | -1.166 | L:K31R | -0.944 | L:N35W | -0.773 |
| L:A19D | -1.164 | L:S32F | -0.939 | L:S33K | -0.768 |
| L:A51N | -1.149 | L:G29W | -0.923 | L:S33R | -0.709 |
| L:S52Y | -1.104 | L:S93D | -0.906 | L:S62I | -0.695 |
| H:N57Q | -1.104 | L:G29P | -0.841 | L:I34Y | -0.690 |
| H:A33M | -1.002 | L:N27W | -0.835 | L:S32N | -0.677 |
| H:S31F | -0.995 | L:N27Y | -0.822 | L:S62Q | -0.673 |

### Structural inspection

During the structural inspection, physiochemical factors, such as spatial limitations, removal of salt bridge or hydrogen bond, deletion or introduction of Cys, Met, and Pro residues were taken into consideration. As shown in Figure 2C and Table 1, selected mutations for 80R belong to positions D50, A51, S52, S67, S92 in the light chain; and N57, R100, S101 in the heavy chain. Since A51 is in the vicinity of Y489, F490, and Q493, it is expected that mutations to Phe, Trp, or Tyr will promote formation of π—π interactions, while mutations to Glu, His, Arg, and Lys may facilitate hydrogen bond interactions with Q429. Similarly, mutations at site 50 and 32 can either strengthen the hydrogen bond or form nonpolar interactions with the bonding partners on the RBD. Side chains of residue 100 and 101 on the heavy chain are in close proximity to Y505, therefore, introduction of aromatic side chains in these locations are presumably favorable. Heavy chain S101D mutation was selected due to possible hydrogen bond enhancement for interacting with N501 (Figure 4C).

As shown in Figure 3C and Table 1, top ranked affinity enhancing mutations for m396 are primarily located at the CDRH2 loop, such as residues 52—59. These residues are in a close contact with R403, Q498, Y505, and N501 on the RBD. H:I57R and H:N59R mutations can introduce a salt bridge with D405 resulting in stronger binding to the RBD. H:S31X mutations, where X is polar side chain, increases the possibility of hydrogen bond formation with T500 and N501 on the RBD. Structural investigation does not support the H:S31F change as it disrupts the hydrogen bond network at this site. However, due to the large Z-score and consistency of the three methods, including MOE, Flex ddg, and TopNetTree, this mutation was included in the suggested list. Mutations on the light chain, including L:G29X and L:S30X, where X is an aromatic mutation, is highly favorable as these side chains are in proximity of Y369 and F374. L:S30E, L:S30H, and L:S30K can result in strong hydrogen bond interactions with the backbone of the RBD near L:S30. Lastly, L:S93E may introduce a salt bridge with the R408 side chain. As shown in Figure 4C and Table 1, selected mutations on the light chain of CR3022 are located on 4 sites, 33—35 and 62. The polar substitutions of these residues are justified through possibility of formation of a hydrogen bond network with D428 and T430 on the RBD, whereas nonpolar mutations can enhance the hydrophobic interactions with L517. The selected mutations on H:G101 and H:S103 of the heavy chain are all of aromatic nature due to their proximity to Y380 and F377. Chain elongation and a more polar headgroup in the H:S100Q substitution can potentially enhance the hydrogen bond network with S383, T385, and K386. H:I102Y is likely to enhance interactions with Y380, while H:T104E and H:Y27R mutations could promote a stronger hydrogen bond network with S383 and N370, respectively.

### Developability engineering

Computational developability risk assessments were focused on chemical liability sites that are nearby or within the paratope and surface patch forming residues, such as hydrophobic and charged residues.

De-risk plan for antibodies 80R and CR3022 is proposed only for chemical liabilities. In 80R, CDRH2 largely contributes to RBD binding. H:D54-G55 which sit in the middle of CDRH2 are considered high risk, although H:D54 does not directly contact RBD residues; H:D54E mutation is therefore proposed. In CR3022, D54 in the DS motif in CDRH2 forms salt bridge to K378 in the RBD. H:D54E mutation is proposed, as H:D54 flanking residues are not directly interacting with the RBD and H:D54 sits in a relatively flexible loop. Surface patches on those two antibodies are generally smaller than 100 Å^2^ and are considered as low risk. One cluster of hydrophobic residues exists in CR3022 around CDRL2 and the 17-residue long CDRL1. The hydrophobic interface (L-I34, Y55, W56) is critical for RBD binding, and the surface is surrounded by charged residues. Therefore, no mitigation plan is proposed on the CR3022 hydrophobic patch.

Among the three selected antibodies, m396 has the highest developability risk, with a 130 Å^2^hydrophobic patch around CDRH2 (H:I54-L55-G56-I57), and a 130 Å^2^acidic patch around CDRL2 (L:D50-D51-S52-D53) (Figure S1B). Chemical liabilities in m396, including exposed H:M102 in CDRH3, L:N26-N27 motif in CDRL1 and L:D92-S93 motif in CDRL3, were predicted as moderate risk since those residues are not directly mediating RBD recognition. To mitigate the risks in m396, mutations giving higher consensus scores at residues L:N26, L:D51, L:D92, H:I54, H:L55, H:I57, and H:M102 were selected into the screening library (Figure 3D, Tables 1, 3).

**Table 3.** Selected m396 mutations for library design. Wild-type residues are listed in bold. Underlined residues are potential developability labile sites.

|  | m396: variable light chain |  |  |  |  |  |  |  |  |  | m396: variable heavy chain |  |  |  |  |  |  |  |  |  |
| --- | --- | --- | --- | --- | --- | --- | --- | --- | --- | --- | --- | --- | --- | --- | --- | --- | --- | --- | --- | --- |
|  | 26 | 27 | 29 | 30 | 31 | 32 | 51 | 92 | 93 | 94 | 95 | 31 | 52 | 54 | 55 | 57 | 59 | 101 | 102 | 103 |
| wild type | <u>N</u> | **N** | **G** | **S** | **K** | **S** | <u>D</u> | <u>D</u> | **S** | **S** | **S** | **S** | **T** | <u>I</u> | <u>L</u> | <u>I</u> | **N** | **V** | <u>M</u> | **G** |
|  | Q | E | F | Y | R | W | W | E | Y | F | I | R | Y | F | Y | R | F | Y | W | F |
|  |  | W | R | F |  | F |  |  | I | W |  | F |  |  | F | W | R | W | Y | W |
|  |  | Y | W | W |  |  |  |  | N |  |  | Y |  |  |  | H | Y | F |  |  |
|  |  |  | P | R |  |  |  |  | D |  |  | Q |  |  |  |  |  | L |  |  |
|  |  |  |  | H |  |  |  |  |  |  |  | W |  |  |  |  |  | R |  |  |
|  |  |  |  | E |  |  |  |  |  |  |  | K |  |  |  |  |  |  |  |  |
|  |  |  |  | K |  |  |  |  |  |  |  |  |  |  |  |  |  |  |  |  |

### Library design

With proposed affinity and developability optimization mutations, we next proceeded to design three focused libraries for 80R, m396, and CR3022 individually. The designed libraries will be used by a high-throughput system, such as phage display, to screen for high affinity binders. Table 2-4 summarizes variations at different positions in the three libraries. The resulting theoretical library sizes are all smaller than 1 × 10^11^, which are suitable for phage display screening.

**Table 2.**
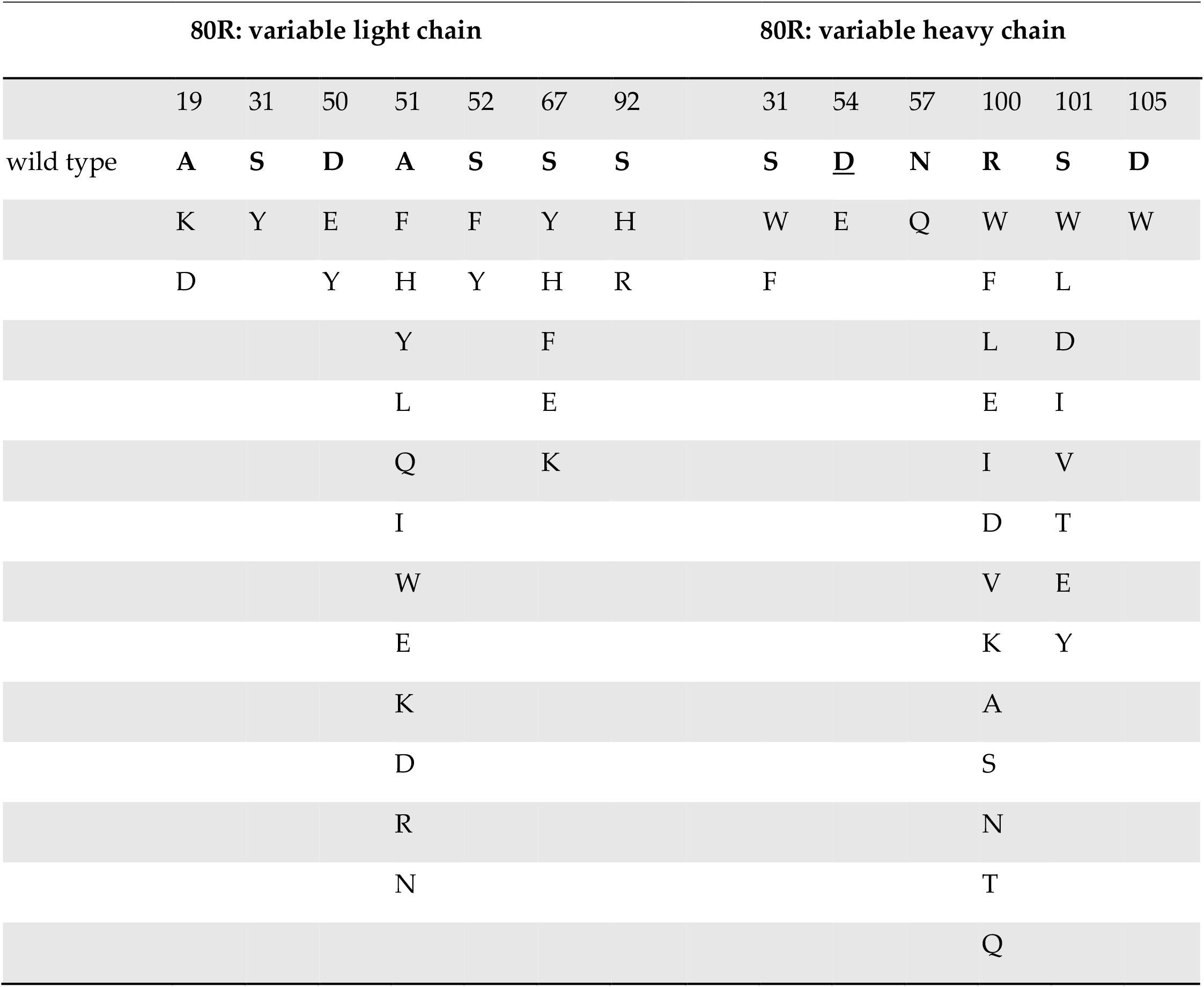
Selected 80R mutations for library design. Wild-type residues are listed in bold. Underlined residues are potential developability labile sites.

**Table 4.**
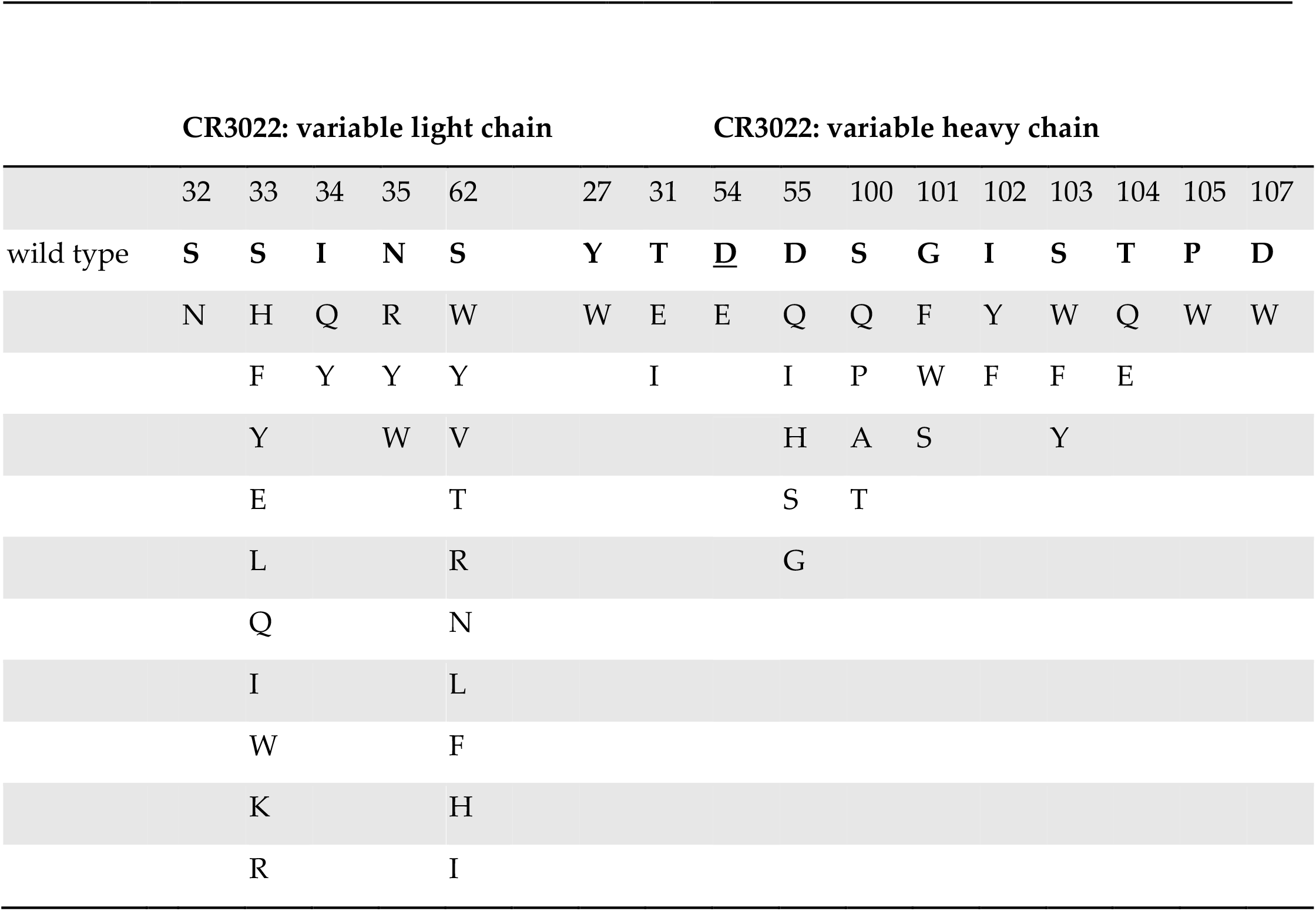
Selected CR3022 mutations for library design. Wild-type residues are listed in bold. Underlined residues are potential developability labile sites.

### Trispecific antibody design and Fc selection

SARS-CoV-2 has shown fast mutation rates among discovered variants, therefore combining neutralizing antibodies with different epitopes into a multi-specific format can benefit both potency and breadth, especially for future variants. We therefore proposed to engineer the three mAbs, after affinity optimization against SARS-CoV-2, into a trispecific format, which has been demonstrated successful in HIV neutralization [14]. The trispecific format includes a single Fab arm derived from a normal immunoglobulin G (IgG) with a double Fv arm generated in the CODV-Ig format (cross-over dual variable Ig-like proteins) [46] (Figure 5A-B). We modeled all possible combinatorial structures of CODV in complex with SARS-CoV-2 spike proteins (Figure 5C-D). Interestingly, it had been reported that CR3022 binding requires rearrangements in the S1 domain of the spike protein which results in dissociation of the spike [47]. A similar observation that CR3022 showed incompatibility to all possible CODV configurations led us to keep CR3022 in the Fab arm and use m396 and 80R in the CODV arm. After examining the structural compatibility, option 2 (80R as VH1/VL2, m396 as VH2/VL1) showed to be the best geometrical configuration (Figure 5C-D).

**Figure 5.**
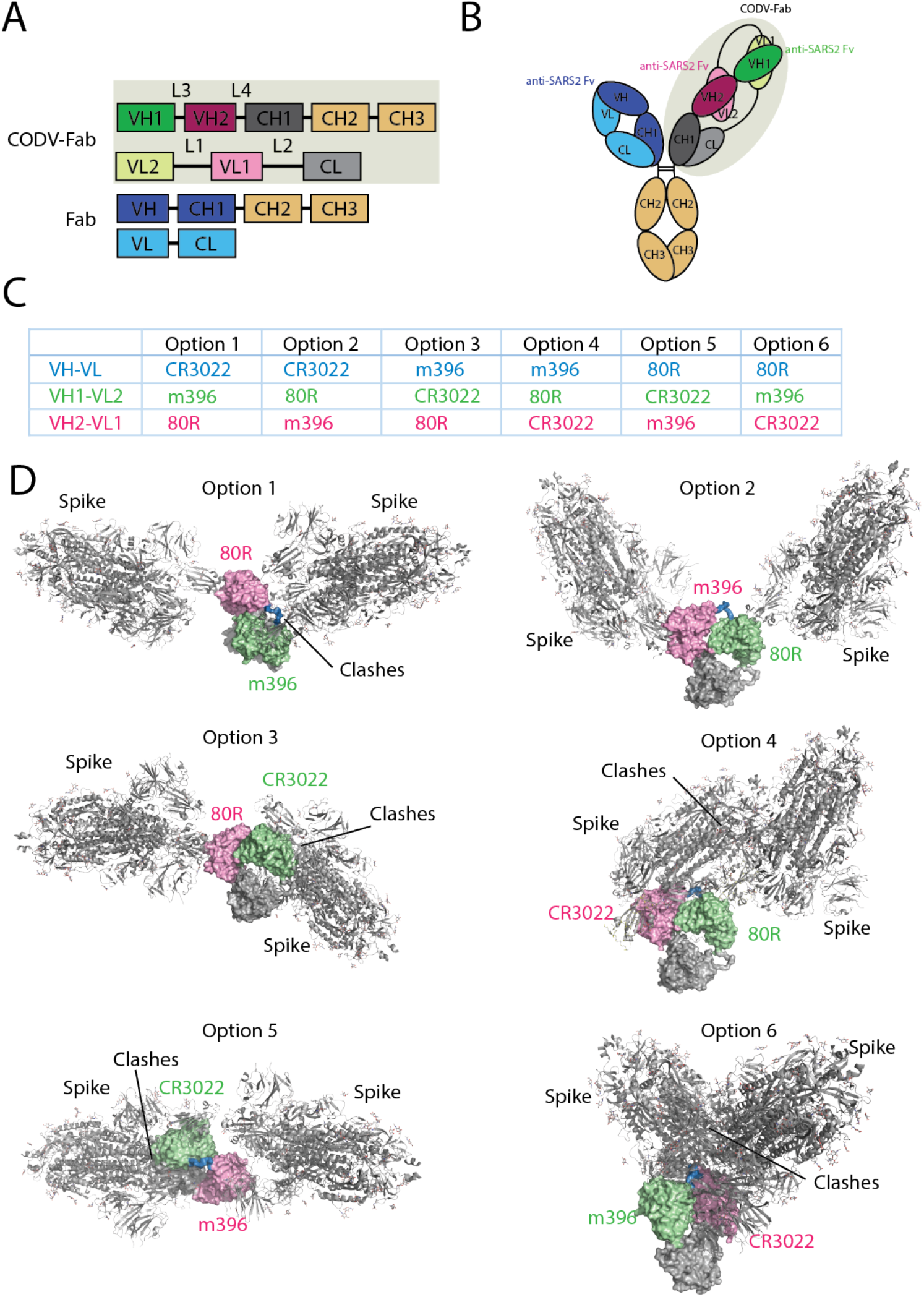
Trispecific antibody engineering. **(A)** Schematic linear configuration of the trispecific antibody color-coded by position. Dark shades (blue, purple, or green) denote heavy chain peptides; light shades denote light chain peptides. **(B)** Schematic cartoon configuration of the trispecific antibody shown in cartoon. Same color scheme is used as that in (A). **(C)** All possible combinations of the three Fvs in the trispecific format. **(D)** Structural modeling showed only Option 2 as the optimal geometrical configuration. The CODV is shown in surface format and color coded as in (A-C), and the spike proteins are shown in grey colored cartoon.

Modifications to the Fc domain are devised to block the contact formation between the Fc region and effector cells. Antibody-dependent enhancement (ADE) potentially poses a safety risk to an antibody treatment, and anti-SARS-CoV-2 antibodies could exacerbate COVID-19 through antibody-dependent enhancement [48]. Although effector function has been recently reported as essential for optimal efficacy in SARS-CoV-2 monoclonal antibody SC31 [49], considering the triplicated valency in our CODV-IgG trispecific antibody, we included NNAS glycosylation at the FcγR interface [50] to completely eliminate Fc-mediated effector functions therefore minimizing ADE risk, and DQ mutations at the FcRn interface [51] to extend antibody half-life.

## Discussion

The devastating COVID-19 pandemic urges faster and smarter designs of treatment to patients worldwide. Antibody therapies have been shown to have the advantages of large-scale production and anti-viral potency. Structure-based rational engineering to redesign well characterized SARS-CoV neutralizing mAbs enables quick solutions to create a pool of SARS-CoV-2 neutralizers with known epitopes. In this work we share our knowledge in antibody engineering especially in multi-specific formats. Using computational protein engineering tools, we proposed a multi-specific antibody based on optimization of SARS-CoV neutralizing antibodies. Our extensive exploration of mutational space involved in the direct interaction with the SARS-CoV-2 RBD has produced a mutation library that is expected to improve the efficacy of these antibodies against the SARS-CoV-2 virus. Physiochemical properties and free energy calculations of each mutation were taken into consideration in building our mutation library. The satisfactory level of agreement and consistency among three of the methods used in this study, including MOE, Rosetta Flex ddg, and TopNetTree, highlights the effectiveness of our proposed library design.

Several AI-guided studies have been carried out to discover treatment against SARS-CoV-2 virus, including the work from Magar et al. [52] and Desautels et al. [53]. Using a ML-based algorithm, Magar and coworkers proposed single and combinatorial mutations on 80R and S230 antibodies with potentially better antibody response. In the case of 80R, the proposed mutations are largely distal from the binding site, and they don’t overlap with our proposal. Since the ML-based model was trained on patient neutralization response, it may capture different properties related to neutralization rather than direct interaction with antigen. It is intriguing that there may be a synergistic effect when combining the ML-based mutations with our proposed mutations in neutralization activity. In another work from Desautels et al., antibody candidates were proposed using an active learning protocol where the model takes Rosetta scores as ground truth and continuously improves its predictability. Complex structures of SARS-CoV neutralizing Abs, including S230, m396, and F26G19, were fed to the algorithm, and mutants with favorable predicted Rosetta scores were proposed. The mutants were further selected by free energy calculations using MD simulations under the implicit solvent model (GBSA). After all, mutations were selected based on Rosetta score and MM/GBSA free energy, while the ML model was used to predict Rosetta scores of large numbers of mutations. In contrast, our method used two separate ML-based models predicting affinity changes directly and assembled the results together with two physics-based methods, one Rosetta-based and one similar to MM/GBSA. By this more diverse scoring system, we expect to increase the prediction accuracy. Moreover, a high-throughput screening method enables testing more mutations and their combinations, which will further increase the possibility of success.

Continuous evolution of SARS-CoV-2 virus remains a significant threat even after the successes of current vaccine development. Among the mutations in the UK and South African strains, E484K is within the 80R epitope, while N501Y is within both 80R and m396 epitopes (Figure 1B). This emphasizes the importance of combining multiple antibodies with different epitopes, especially to include antibodies with conserved epitopes, such as CR3022. Given the success shown in the HIV study, our trispecific format is one of the suitable formats for 3-in-1 antibody design. However, it requires careful geometry modeling and sequence optimization for further developability.

## Conclusions

In this study, we used computational protein engineering tools to optimize SARS-CoV neutralizing mAbs against SARS-CoV-2 virus. Three mAbs were used as templates where their complex structures with SARS-CoV-2 RBD were optimized following modeling protocols in Rosetta and MOE simulation packages. Subsequently, extensive free energy calculations were carried out on the residues in contact with the RBD. Two physics-based and two ML-based free energy calculation suites were utilized to perform the affinity maturation calculations. For each system, developability assessment was done and a focused library was proposed for high-throughput screening of high affinity and developable Fabs against the SARS-CoV-2 RBD. Lastly, a design of combining the three antibodies in a trispecific format was achieved, aiming for high potency and broad neutralization activity.

**Figure S1.**
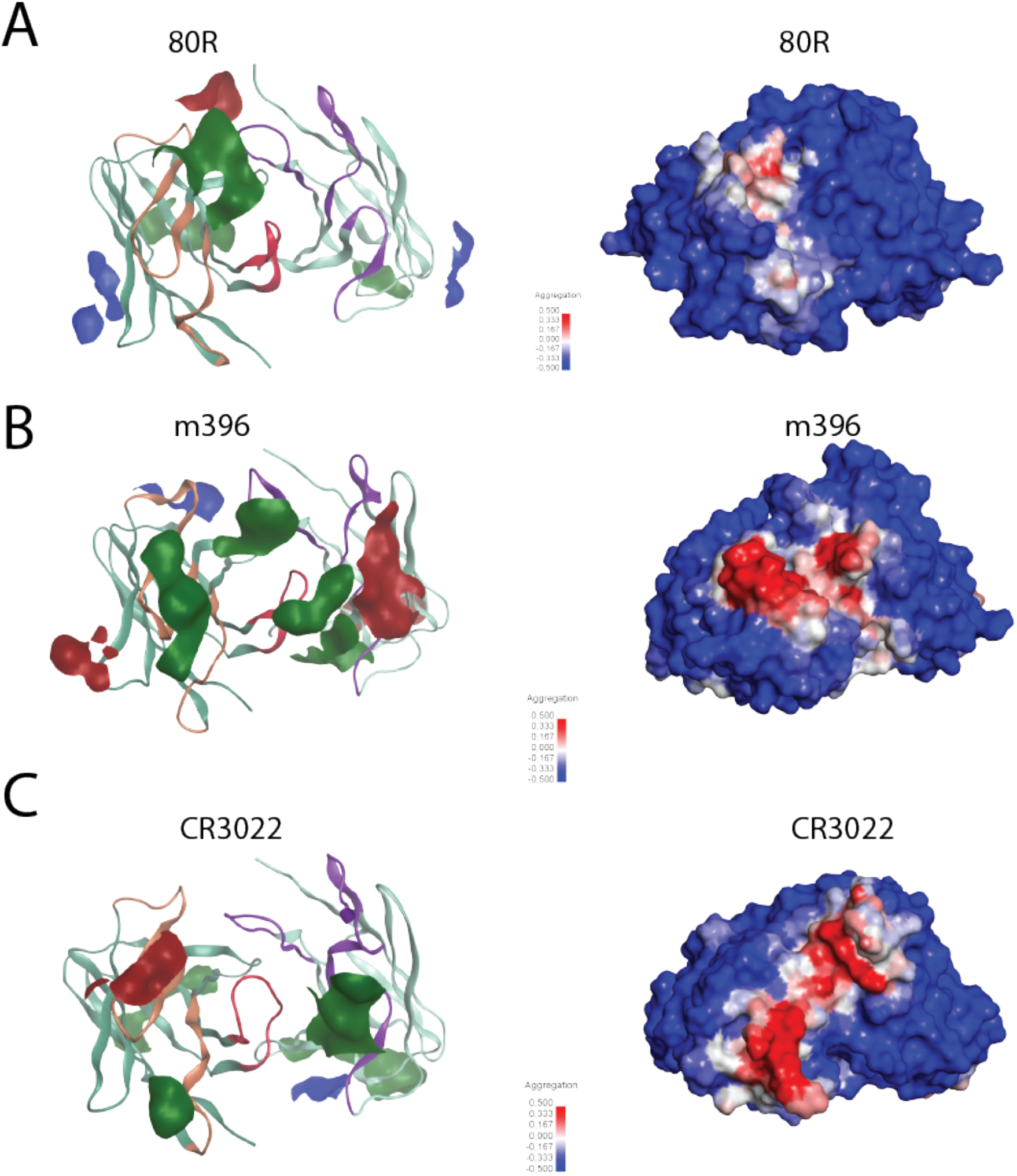
Developability assessment on antibody 80R, m396, and CR3022. Left: MOE patch analysis on the Fv region of antibody **(A)** 80R, **(B)** m396, and **(C)** CR3022. Red color indicates negative-charge patch, blue color indicates positive-charge patch, and green color indicates hydrophobic patch. Right: spatial aggregation propensity (SAP) analysis of antibody **(A)** 80R, **(B)** m396, and **(C)** CR3022, with high SAP score colored in red and low SAP score colored in blue (scale showed in a bar scheme).

